# A vector system for fast-forward *in vivo* studies of the ZAR1 resistosome in the model plant *Nicotiana benthamiana*

**DOI:** 10.1101/2020.05.15.097584

**Authors:** Adeline Harant, Toshiyuki Sakai, Sophien Kamoun, Hiroaki Adachi

## Abstract

*Nicotiana benthamiana* has emerged as a complementary experimental system to Arabidopsis. It enables fast-forward *in vivo* analyses primarily through transient gene expression and is particularly popular in the study of plant immunity. Recently, our understanding of NLR plant immune receptors has greatly advanced following the discovery of Arabidopsis ZAR1 resistosome. Here, we describe a novel vector system of 52 plasmids that enables functional studies of the ZAR1 resistosome in *N. benthamiana*. We showed that ZAR1 stands out among the coiled coil class of NLRs for being highly conserved across distantly related dicot plant species and confirmed NbZAR1 as the *N. benthamiana* ortholog of Arabidopsis ZAR1. NbZAR1 triggers autoimmune cell death in *N. benthamiana* and this activity is dependent on a functional N-terminal α1 helix. C-terminally tagged NbZAR1 remains functional in *N. benthamiana* thus enabling cell biology and biochemical studies in this plant system. We conclude that the NbZAR1 open source plasmids form an additional experimental system to Arabidopsis for *in planta* resistosome studies.

## INTRODUCTION

*Nicotiana benthamiana* has developed into a popular model system in plant biology, particularly for the study of plant immunity. The appeal of *N. benthamiana* as an experimental system is primarily due to agroinfiltration, a rapid transient protein expression method that enables cell biology, biochemistry, protein–protein interaction analyses and other *in vivo* experiments (Goodin et al., 2008). In addition, *N. benthamiana* is a member of the asterid group of flowering plants and as such is relatively distant from the classic model plant *Arabidopsis thaliana*, which belongs to the rosid group. Therefore, *N. benthamiana* is a complementary experimental system to Arabidopsis that also allows for a broader perspective into the evolution of molecular mechanisms in dicot plants.

Plants use nucleotide-binding leucine-rich repeat (NLR) proteins to mount innate immunity against invading pathogens. NLRs are intracellular receptors that detect host cell-translocated pathogen effectors and activate a hypersensitive cell death response that is a hallmark of plant immunity. The recent elucidation of the structures of the inactive and active complexes formed by the Arabidopsis NLR protein ZAR1 (HOPZ-ACTIVATED RESISTANCE1) with its partner receptor-like cytoplasmic kinases (RLCKs) is a major breakthrough in understanding how these immune receptors are activated and function. Activated ZAR1 forms a wheel-like pentamer, termed resistosome, that undergoes a conformational switch—the death switch— to expose a funnel-shaped structure formed by the N-terminal α1 helices (Wang et al., 2019a; 2019b). The ZAR1 resistosome was proposed to mediate cell death directly by translocating into the plasma membrane through the funnel-like structure and perturbing membrane integrity similar to pore-forming toxins (Wang et al., 2019b). The N-terminal α1 helix matches a sequence motif known as MADA motif that is functionally conserved across ∼20% of coiled-coil (CC)-type plant NLRs (Adachi et al., 2019). This indicates that the ZAR1 death switch mechanism may be widely conserved across plant species. Mutations in surface-exposed residues within the α1 helix/MADA motif impair cell death and disease resistance mediated by ZAR1 and other MADA-CC-NLRs (Wang et al., 2019b; Adachi et al., 2019).

Here, we describe a Golden Gate compatible vector system (pZA) that enables functional studies of the ZAR1 resistosome in *N. benthamiana*. Golden Gate enables modular cloning through assembly of DNA fragments in one-step Type IIS restriction enzyme reactions. Consistent with previous reports (Baudin et al. 2017; 2019), we confirmed that Arabidopsis ZAR1 (AtZAR1) does not cause hypersensitive cell death in *N. benthamiana* when expressed on its own and that *N. benthamiana* carries an ortholog of ZAR1 (NbZAR1). We developed and validated the pZA plasmid collection and share it as an open source resource that should prove useful for functional and comparative studies of NLR resistosomes in plants.

## RESULTS AND DISCUSSION

### ZAR1 is conserved across distantly related dicot plant species

Baudin et al. (2017) previously reported two orthologs of Arabidopsis ZAR1 and named them NbZAR1 and NbZAR2. We confirmed their findings by performing phylogenetic analyses of NLR proteins using the NB-ARC (nucleotide-binding adaptor shared by APAF-1, certain *R* gene products and CED-4) domain from 5 dicot plant species (Arabidopsis, cassava, sugar beet, tomato and *N. benthamiana*) representing three major clades of dicot plants (rosids, aterids and Caryophyllales). The resulting phylogenetic tree included 829 NLRs, which expectedly clustered into the well-defined NLR classes (Figure 1A). Arabidopsis ZAR1 (AtZAR1, AT3G50950) fell within a well-supported CC-NLR subclade that included NbZAR1 (NbS00000462g0011), NbZAR2 (NbS00013646g0007) and one gene each from the three other dicot plant species (Figure 1A). As previously reported (Baudin et al. 2017), NbZAR2 gene is truncated in the reference *N. benthamiana* genome (Figure 1—figure supplement 1). The other four genes, NbZAR1, Manes.02G097400 (MeZAR1), Bv9 203570 geqf.t1 (BvZAR1) and Solyc02g084890 (SlZAR1), code for CC-NLR proteins of similar length to AtZAR1, suggesting them as ZAR1 orthologs in each dicot plant species (Figure 1—figure supplement 1).

**Figure 1.**
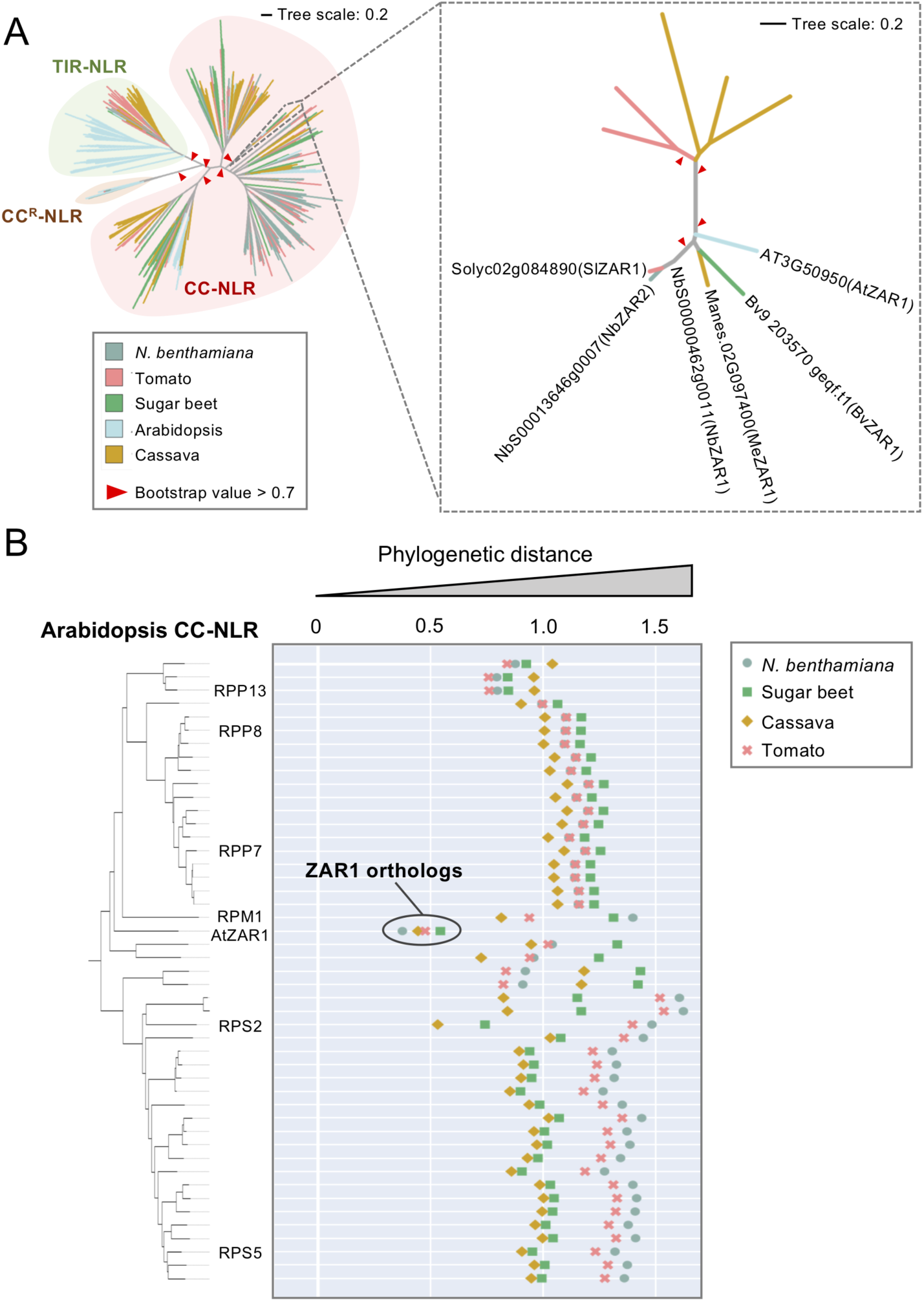
ZAR1 is a conserved CC-NLR across distantly related dicot plant species. (**A**) ZAR1 forms a small subclade that is composed of orthologous genes from distantly related dicot plant species. The phylogenetic tree was generated in MEGA7 by the neighbour-joining method using NB-ARC domain sequences of 829 NLRs identified from *N. benthamiana* (NbS-), tomato (Solyc-), sugar beet (Bv-), Arabidopsis (AT-) and cassava (Manes-) (left panel). The ZAR1 subclade phylogenetic tree is shown on the right panel. Each branch is marked with different colours based on plant species. Red arrow heads indicate bootstrap support > 0.7 and is shown for the relevant nodes. The scale bars indicate the evolutionary distance in amino acid substitution per site. Amino acid sequences of the full-length ZAR1 orthologs can be found in Figure 1—figure supplement 1. (**B**) AtZAR1 is highly conserved among dicot CC-NLRs. The phylogenetic (patristic) distance of two CC-NLR nodes between Arabidopsis and other plant species were calculated from the NB-ARC phylogenetic tree in A. The closest patristic distances are plotted with different colours based on plant species. Representative Arabidopsis NLRs are highlighted. The closest patristic distances of two CC-NLR nodes between *N. benthamiana* and other plant species can be found in Figure 1—figure supplement 2.

Based on our experience, ZAR1 seemed atypical among NLRs for having well-defined orthologs across unrelated dicot species. To directly evaluate ZAR1 conservation relative to other CC-NLRs, we calculated the phylogenetic (patristic) distance between each of the 47 Arabidopsis CC-NLRs and their closest gene from the other plant species based on the phylogenetic tree (Figure 1A). Interestingly, AtZAR1 stood out among the CC-NLRs as having the shortest phylogenetic distance to its four orthologs (Figure 1B). Similarly, in a reverse analysis where we plotted the phylogenetic distance between each of the 153 *N. benthamiana* CC-NLRs to their closest gene from the other species, NbZAR1 again stood out as being exceptionally conserved across all five examined species relative to other *N. benthamiana* CC-NLRs (Figure 1—figure supplement 2). Multiple alignments of the ZAR1 orthologs from the examined species confirmed the relatively high conservation of ZAR1 with pair-wise sequence similarities ranging from 58% to 86% across the full NLR protein (Figure 1—figure supplement 1). Overall, these analyses confirm NbZAR1 as the *N. benthamiana* ortholog of AtZAR1 and reveal ZAR1 as possibly the most widely conserved CC-NLR in dicot plants.

### Golden Gate compatible ZAR1 plasmids for *in vivo* resistosome studies

The Golden Gate cloning system enables rapid and high throughput assembly of multiple sequence modules, such as promotor, terminator or tags into a common vector (Patron et al., 2015). To facilitate *in vivo* resistosome studies using this cloning strategy, we first introduced synonymous mutations to *Bpi*I restriction enzyme sites in the coding sequence of *AtZAR1* and *NbZAR1* to make them both Golden Gate compatible and enable their transfer into a variety of *Agrobacterium tumefaciens* binary vectors for *in planta* protein expression. We cloned the genes into the Golden Gate compatible vector pUC8-Spec (pZA1, 5, 9 and 13 in Table 1). Each coding sequence was flanked by *Bsa*I restriction enzyme sites and overhang sequences for optimal Golden Gate reactions (Table 1—figure supplement 1).

**Table 1.** Golden Gate Plasmid resources prepared in this study.

| Plasmids | Relevant features | Bacterial selection | Usage |
| --- | --- | --- | --- |
| pZA1 | Golden Gate compatible (GG_) wild-type AtZAR1 with STOP codon | Spc<sup>r</sup> | Level 0 |
| pZA2 | GG_AtZAR1<sup>D489V</sup> with STOP codon | Spc<sup>r</sup> | Level 0 |
| pZA3 | GG_AtZAR1<sup>L17E</sup> with STOP codon | Spc<sup>r</sup> | Level 0 |
| pZA4 | GG_AtZAR1<sup>L17E/D489V</sup> with STOP codon | Spc<sup>r</sup> | Level 0 |
| pZA5 | GG_wild-type NbZAR1 with STOP codon | Spc<sup>r</sup> | Level 0 |
| pZA6 | GG_NbZAR1<sup>D481V</sup> with STOP codon | Spc<sup>r</sup> | Level 0 |
| pZA7 | GG_NbZAR1<sup>L17E</sup> with STOP codon | Spc<sup>r</sup> | Level 0 |
| pZA8 | GG_NbZAR1<sup>L17E/D481V</sup> with STOP codon | Spc<sup>r</sup> | Level 0 |
| pZA9 | GG_wild-type AtZAR1 without STOP codon | Spc<sup>r</sup> | Level 0 |
| pZA10 | GG_AtZAR1<sup>D489V</sup> without STOP codon | Spc<sup>r</sup> | Level 0 |
| pZA11 | GG_AtZAR1<sup>L17E</sup> without STOP codon | Spc<sup>r</sup> | Level 0 |
| pZA12 | GG_AtZAR1<sup>L17E/D489V</sup> without STOP codon | Spc<sup>r</sup> | Level 0 |
| pZA13 | GG_wild-type NbZAR1 without STOP codon | Spc<sup>r</sup> | Level 0 |
| pZA14 | GG_NbZAR1<sup>D481V</sup> without STOP codon | Spc<sup>r</sup> | Level 0 |
| pZA15 | GG_NbZAR1<sup>L17E</sup> without STOP codon | Spc<sup>r</sup> | Level 0 |
| pZA16 | GG_NbZAR1<sup>L17E/D481V</sup> without STOP codon | Spc<sup>r</sup> | Level 0 |
| pZA17 | GG_wild-type AtZAR1, Km<sup>r</sup> plant selection marker | Km<sup>r</sup> | Level 2 |
| pZA18 | GG_AtZAR1<sup>D489V</sup>, Km<sup>r</sup> plant selection marker | Km<sup>r</sup> | Level 2 |
| pZA19 | GG_AtZAR1<sup>L17E</sup>, Km<sup>r</sup> plant selection marker | Km<sup>r</sup> | Level 2 |
| pZA20 | GG_AtZAR1<sup>L17E/D489V</sup>, Km<sup>r</sup> plant selection marker | Km<sup>r</sup> | Level 2 |
| pZA25 | GG_wild-type NbZAR1, Km<sup>r</sup> plant selection marker | Km<sup>r</sup> | Level 2 |
| pZA26 | GG_NbZAR1<sup>D481V</sup>, Km<sup>r</sup> plant selection marker | Km<sup>r</sup> | Level 2 |
| pZA27 | GG_NbZAR1<sup>L17E</sup>, Km<sup>r</sup> plant selection marker | Km<sup>r</sup> | Level 2 |
| pZA28 | GG_NbZAR1<sup>L17E/D481V</sup>, Km<sup>r</sup> plant selection marker | Km<sup>r</sup> | Level 2 |

**Table 1.** Continued.
| Plasmids | Relevant features | Bacterial selection | Usage |
| --- | --- | --- | --- |
| pZA29 | GG_wild-type AtZAR1-4xMyc, Km<sup>r</sup> plant selection marker | Km<sup>r</sup> | Level 2 |
| pZA30 | GG_AtZAR1<sup>D489V</sup>-4xMyc, Km<sup>r</sup> plant selection marker | Km<sup>r</sup> | Level 2 |
| pZA31 | GG_AtZAR1<sup>L17E</sup>-4xMyc, Km<sup>r</sup> plant selection marker | Km<sup>r</sup> | Level 2 |
| pZA32 | GG_AtZAR1<sup>L17E/D489V</sup>-4xMyc, Km<sup>r</sup> plant selection marker | Km<sup>r</sup> | Level 2 |
| pZA33 | GG_wild-type NbZAR1-4xMyc, Km<sup>r</sup> plant selection marker | Km<sup>r</sup> | Level 2 |
| pZA34 | GG_NbZAR1<sup>D481V</sup>-4xMyc, Km<sup>r</sup> plant selection marker | Km<sup>r</sup> | Level 2 |
| pZA35 | GG_NbZAR1<sup>L17E</sup>-4xMyc, Km<sup>r</sup> plant selection marker | Km<sup>r</sup> | Level 2 |
| pZA36 | GG_NbZAR1<sup>L17E/D481V</sup>-4xMyc, Km<sup>r</sup> plant selection marker | Km<sup>r</sup> | Level 2 |
| pZA37 | GG_wild-type AtZAR1-6xHA, Km<sup>r</sup> plant selection marker | Km<sup>r</sup> | Level 2 |
| pZA38 | GG_AtZAR1<sup>D489V</sup>-6xHA, Km<sup>r</sup> plant selection marker | Km<sup>r</sup> | Level 2 |
| pZA39 | GG_AtZAR1<sup>L17E</sup>-6xHA, Km<sup>r</sup> plant selection marker | Km<sup>r</sup> | Level 2 |
| pZA40 | GG_AtZAR1<sup>L17E/D489V</sup>-6xHA, Km<sup>r</sup> plant selection marker | Km<sup>r</sup> | Level 2 |
| pZA41 | GG_wild-type NbZAR1-6xHA, Km<sup>r</sup> plant selection marker | Km<sup>r</sup> | Level 2 |
| pZA42 | GG_NbZAR1<sup>D481V</sup>-6xHA, Km<sup>r</sup> plant selection marker | Km<sup>r</sup> | Level 2 |
| pZA43 | GG_NbZAR1<sup>L17E</sup>-6xHA, Km<sup>r</sup> plant selection marker | Km<sup>r</sup> | Level 2 |
| pZA44 | GG_NbZAR1<sup>L17E/D481V</sup>-6xHA, Km<sup>r</sup> plant selection marker | Km<sup>r</sup> | Level 2 |
| pZA45 | GG_wild-type AtZAR1-eGFP, Km<sup>r</sup> plant selection marker | Km<sup>r</sup> | Level 2 |
| pZA46 | GG_AtZAR1<sup>D489V</sup>-eGFP, Km<sup>r</sup> plant selection marker | Km<sup>r</sup> | Level 2 |
| pZA47 | GG_AtZAR1<sup>L17E</sup>-eGFP, Km<sup>r</sup> plant selection marker | Km<sup>r</sup> | Level 2 |
| pZA48 | GG_AtZAR1<sup>L17E/D489V</sup>-eGFP, Km<sup>r</sup> plant selection marker | Km<sup>r</sup> | Level 2 |
| pZA49 | GG_wild-type NbZAR1-eGFP, Km<sup>r</sup> plant selection marker | Km<sup>r</sup> | Level 2 |
| pZA50 | GG_NbZAR1<sup>D481V</sup>-eGFP, Km<sup>r</sup> plant selection marker | Km<sup>r</sup> | Level 2 |
| pZA51 | GG_NbZAR1<sup>L17E</sup>-eGFP, Km<sup>r</sup> plant selection marker | Km<sup>r</sup> | Level 2 |
| pZA52 | GG_NbZAR1<sup>L17E/D481V</sup>-eGFP, Km<sup>r</sup> plant selection marker | Km<sup>r</sup> | Level 2 |

Spc<sup>r</sup>: spectinomycin-resistant, Km<sup>r</sup>: kanamycin-resistant.

We generated autoactive and inactive versions of full-length ZAR1 by mutating the MHD and MADA motifs, which are both conserved across ZAR1 orthologs (Figure 1—figure supplement 1). Whereas mutation from aspartic acid (D) to valine (V) in the MHD motif generally makes full-length NLRs autoactive (Tameling et al., 2006), mutations in the N-terminal MADA motif impair NLR cell death activity (Wang et al., 2019b; Adachi et al., 2019). We generated a mutant series of AtZAR1 and NbZAR1, namely MHD mutants (AtZAR1^D489V^ and NbZAR1^D481V^), MADA mutants (AtZAR1^L17E^ and NbZAR1^L17E^) and MADA/MHD mutants (AtZAR1^L17E/D489V^ and NbZAR1^L17E/D481V^). We cloned the variant genes into pUC8-Spec (pZA2-4, 6-8, 10-12 and 14-16 in Table 1) and assembled the genes into binary vector plasmids without tags or with the C-terminal tags 4xMyc, 6xHA and eGFP (pZA17-52 in Table 1). This collection of 52 plasmids forms an open access resource for *in planta* ZAR1 resistosome studies.

### Unlike Arabidopsis ZAR1, NbZAR1 triggers autoimmune cell death in *N. benthamiana*

To validate the ZAR1 constructs we generated, we first used agroinfiltration to transiently express wild-type ZAR1 and the MHD mutants AtZAR1^D489V^ and NbZAR1^D481V^ in *N. benthamiana* leaves (Figure 2A). As recently reported by Baudin et al. (2019), AtZAR1 did not trigger macroscopic cell death in *N. benthamiana* leaves even when the full-length protein carries the D489V mutation in the MHD motif (Figure 2B and 2C). In contrast, NbZAR1 caused macroscopic cell death when the MHD mutant NbZAR1^D481V^ was expressed in *N. benthamiana* leaves (Figure 2D and 2E). NRC4 was used as a control MADA-CC-NLR, because its MHD mutant NRC4^D478V^ causes autoactive cell death in *N. benthamiana* leaves (Adachi et al., 2019). Our results indicate that unlike Arabidopsis ZAR1, full-length NbZAR1 can be made autoactive in *N. benthamiana* through mutation of the MHD motif. These results are consistent with a previous report that AtZAR1 causes hypersensitive cell death in *N. benthamiana* when co-expressed with the Arabidopsis RLCK AtZED1 (Baudin et al., 2017). We conclude that unlike AtZAR1, NbZAR1 is probably compatible with endogenous RLCKs and can therefore display autoactivity when expressed on its own in *N. benthamiana* leaves.

**Figure 2.**
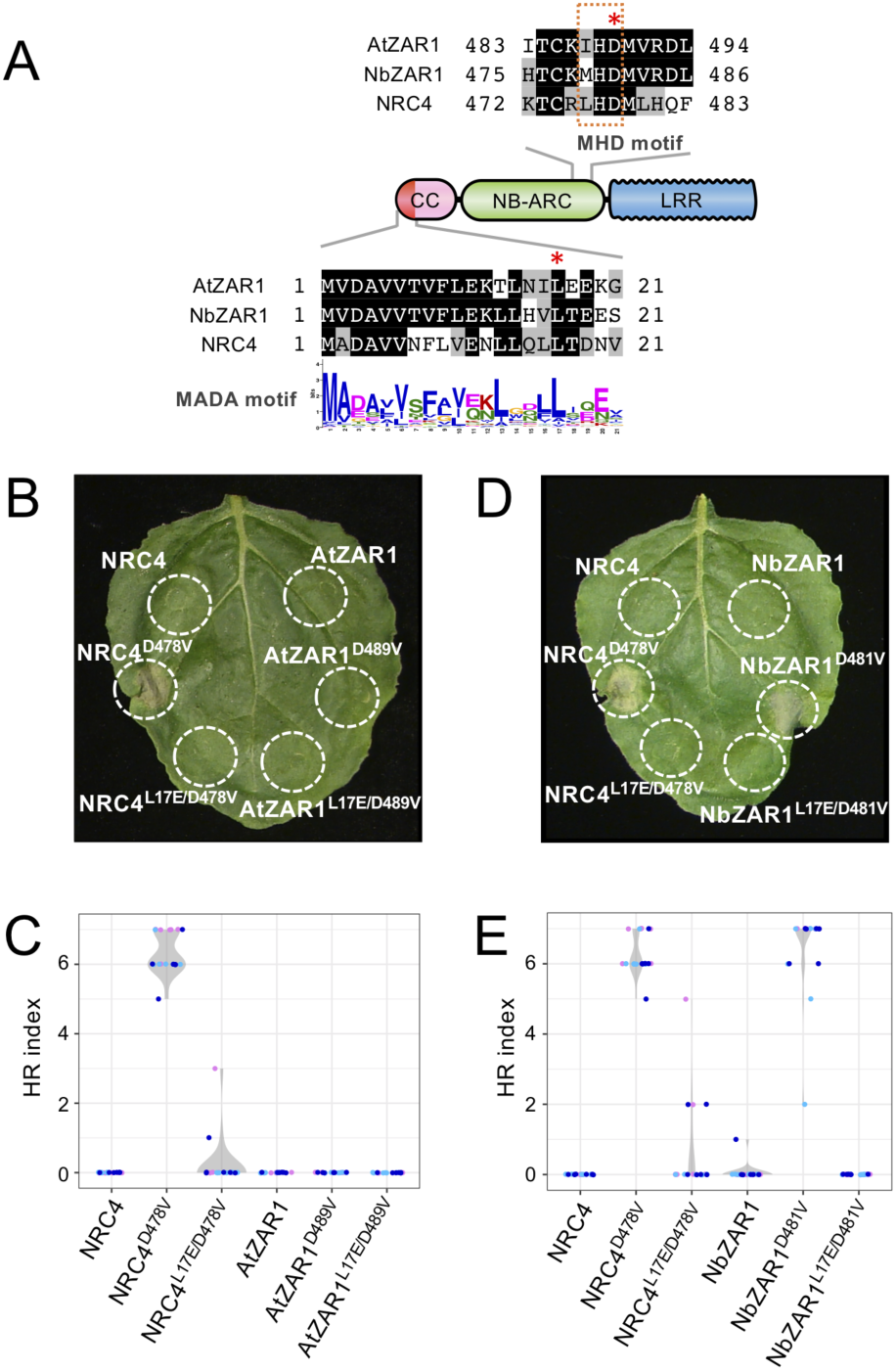
NbZAR1 triggers autoimmune cell death in *N. benthamiana* leaves and requires an intact MADA motif for autoactivity. (**A**) Schematic representation of substitution sites in AtZAR1, NbZAR1 and NRC4. Mutated sites are shown with red asterisks on the sequence alignment. (**B and D**) Cell death observed in *N. benthamiana* leaves after expression of ZAR1 mutants. *N. benthamiana* leaf panels expressing non– epitope-tagged wild-type and variant proteins of AtZAR1 and NbZAR1 were photographed at 5 days after agroinfiltration. (**C and E**) Violin plots showing cell death intensity scored as an HR index based on three independent experiments.

### NbZAR1 autoimmune cell death is dependent on its N-terminal α1 helix

Next, we tested whether NbZAR1 requires an intact N-terminal α1 helix/MADA motif for triggering autoactive cell death. We expressed the double MADA/MHD mutant NbZAR1^L17E/D481V^ in *N. benthamiana* leaves using agroinfiltration and observed that the leucine 17 to glutamic acid mutation significantly reduced the cell death activity compared to NbZAR1^D481V^ (Figure 2D and 2E). This result indicates that an intact N-terminal α1 helix is necessary for NbZAR1^D481V^ autoactivity. Given that the α1 helix forms a funnel-shaped structure that is a key structural component of activated ZAR1 resistosome and its translocation into the plasma membrane (Wang et al., 2019b), we conclude that the NbZAR1^D481V^ experimental system recapitulates the resistosome model in *N. benthamiana*.

### C-terminally tagged NbZAR1 is functional in *N. benthamiana*

*N. benthamiana* agroinfiltration is a powerful method for *in vivo* cell biology, biochemistry and protein–protein interaction analyses, but this experimental system requires that the target protein remains functional and intact after fusion to an epitope or fluorescent tag (Goodin et al., 2008; Bally et al., 2018). We therefore set out to determine the degree to which NbZAR1 can tolerate protein tags (Table 1). We focused on C-terminal tags, given that N-terminal tag fusion is known to interfere with the AtZAR1 cell death activity (Wang et al., 2019b). We tested NbZAR1^D481V^ constructs fused to three different tags, 4xMyc, 6xHA and eGFP by agroinfiltration. In all three cases, NbZAR1^D481V^ retained the capacity to trigger autoactive cell death although the response was weaker than the non–epitope-tagged NbZAR1 positive control (Figure 3A to 3F). We also showed by western blot analyses that these C-terminally tagged NbZAR1 accumulated at detectable levels in *N. benthamiana* leaves and generally formed a single band indicative of minimal protein degradation (Figure 3G to 3I). We conclude that C-terminal tagging of NbZAR1 doesn’t dramatically interfere with the activity and integrity of this NLR protein.

**Figure 3.**
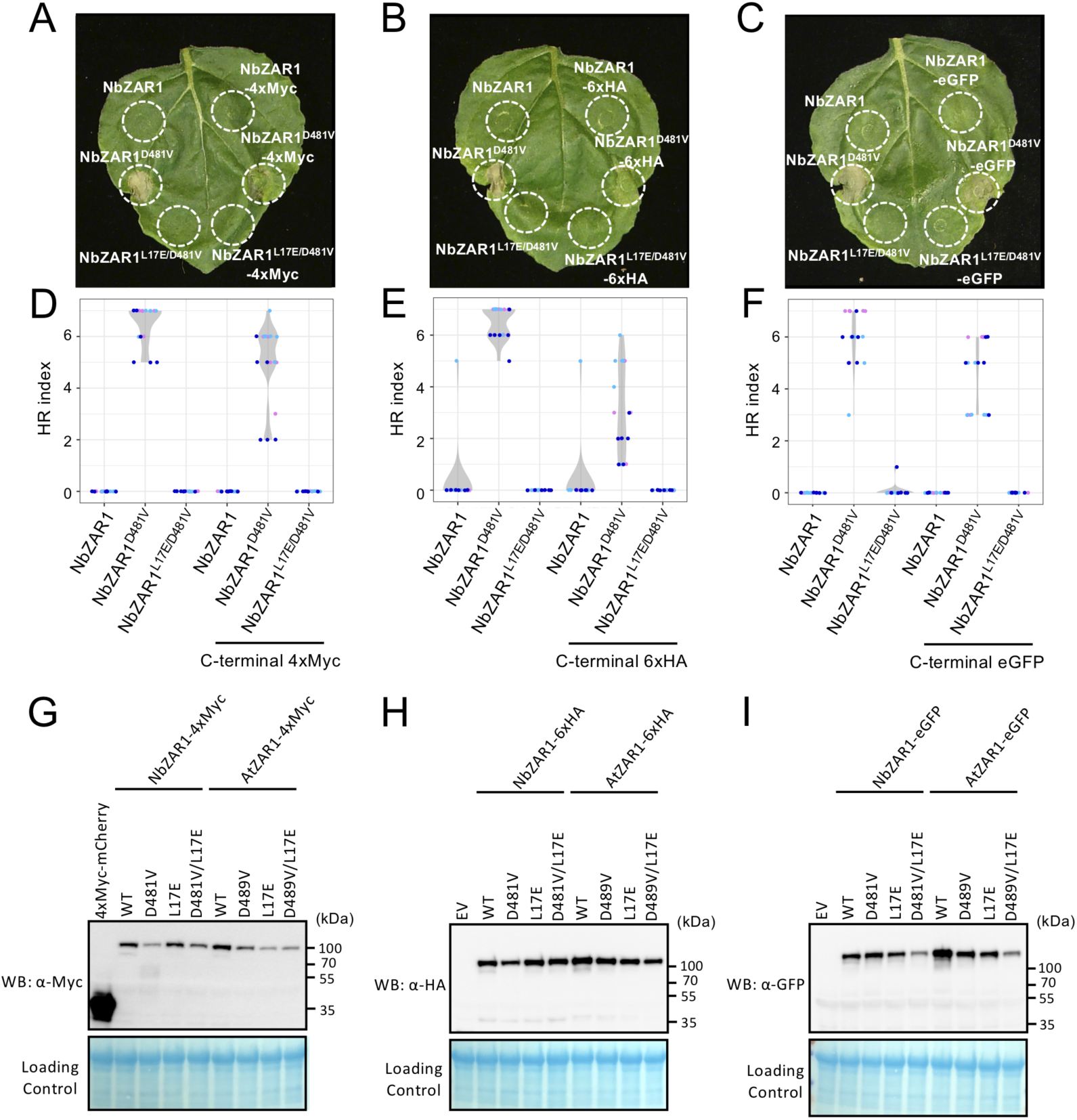
C-terminal tagged NbZAR1 triggers autoimmune cell death in *N. benthamiana* leaves. (**A-C**) Cell death observed in *N. benthamiana* leaves after expression of ZAR1 mutants. *N. benthamiana* leaf panels expressing wild-type and variant proteins of NbZAR1 with C-terminal 4xMyc, 6xHA or eGFP tag were photographed at 5 days after agroinfiltration. (**D-F**) Violin plots showing cell death intensity scored as an HR index based on three independent experiments. (**G-I**) *In planta* protein accumulation of AtZAR1 and NbZAR1 variants. For anti-Myc, anti-HA and anti-GFP immunoblots of AtZAR1, NbZAR1 and the mutant proteins, total proteins were prepared from *N. benthamiana* leaves at 2 days after agroinfiltration. Empty vector control is described as EV. Equal loading was checked with Reversible Protein Stain Kit (Thermo Fisher).

### Towards *in vivo* studies of the ZAR1 resistosome

We conclude that the open source pZA plasmid resource we developed can serve as a valuable experimental system for *in vivo* resistosome studies in *N. benthamiana* that complements the Arabidopsis system and provide a foundation for future studies. The autoimmune NbZAR1^D481V^ mutant can be readily expressed by agroinfilitration in *N. benthamiana* leaves and assayed for cell death activity. NbZAR1^D481V^ tolerates different C-terminal tags enabling *in planta* cell biology, biochemical and protein-protein interactions studies of the ZAR1 resistosome. These tags do not dramatically disturb the activity of NbZAR1^D481V^ and may not affect recruitment of endogenous RLCK partners that are potentially necessary for its function.

The pZA toolkit complements existing assays in Arabidopsis. Of particular interest is the visualization of activated NLR oligomers *in vivo*. Recently, ∼900 kDa complexes associated with activated AtZAR1 and another Arabidopsis MADA-CC-NLR RPP7 were detected using Blue Native polyacrylamide gel electrophoresis and gel filtration (Li et al., 2020; Hu et al., 2020). *N. benthamiana* agroinfiltration offers a rapid and effective alternative expression system to the Arabidopsis protoplast and transgenic expression systems used by Li et al. (2020) and Hu et al. (2020).

ZAR1 stands out among CC-NLRs for being widely conserved across dicot plants but it remains unknown how ZAR1 orthologs vary in their molecular and biochemical activities. The pZA system enables evolutionary studies of the ZAR1 resistosome. Comparative analyses between AtZAR1 and NbZAR1 would help answer questions about the evolution of the resistosome as a mechanism to trigger NLR-mediated immunity.

## MATERIALS AND METHODS

| Key Resources Table |  |  |  |  |
| --- | --- | --- | --- | --- |
| Reagent type (species) or resource | Designation | Source or reference | Identifiers | Additional information |
| Gene (*Arabidopsis thaliana*) | Arabidopsis genome sequence | https://www.araport.org/ | Araport11 | Materials and Methods: Bioinformatic and phylogenetic analyses |
| Gene (*N. benthamiana*) | *N. benthamiana* genome sequence | Sol Genomics Network (https://solgenomics.net/) | *N. benthamiana* Genome v0.4.4 | Materials and Methods: Bioinformatic and phylogenetic analyses |
| Gene (*Solanum lycopersicum*) | Tomato genome sequence | Sol Genomics Network (https://solgenomics.net/) | Tomato ITAG release 2.40 | Materials and Methods: Bioinformatic and phylogenetic analyses |
| Gene (*Manihot esculenta*) | Cassava genome sequence | Phytozome v13 (https://phytozome-next.jgi.doe.gov/) | *Manihot esculenta* v7.1 | Materials and Methods: Bioinformatic and phylogenetic analyses |
| Gene (*Beta vulgaris*) | Sugar beet genome sequence | http://bvseq.molgen.mpg.de/index.shtml | RefBeet-1.2 | Materials and Methods: Bioinformatic and phylogenetic analyses |
| Gene (*Arabidopsis thaliana*) | AtZAR1 | Araport11 (https://www.araport.org/) | AT3G50950 | Materials and Methods: Bioinformatic and phylogenetic analyses |
|  |  | NCBI (https://www.ncbi.nlm.nih.gov/) | NP_190664.1 |  |
| Gene (*N. benthamiana*) | NbZAR1 | N.benthamiana Genome v0.4.4 (https://solgenomics.net/) | NbS00000462g0011 | Materials and Methods: Bioinformatic and phylogenetic analyses |
|  |  | N.benthamiana Genome v1.0.1 (https://solgenomics.net/) | Niben101Scf17398g00012 |  |
|  |  | NCBI (https://www.ncbi.nlm.nih.gov/) | AXY05280.1 |  |
| Gene (*N. benthamiana*) | NbZAR2 | N.benthamiana Genome v0.4.4 (https://solgenomics.net/) | NbS00013646g0007 | Materials and Methods: Bioinformatic and phylogenetic analyses |
|  |  | N.benthamiana Genome v1.0.1 (https://solgenomics.net/) | Niben101Scf00383g03003<br>Niben101Scf00383g03004 |  |
| Gene (*Solanum lycopersicum*) | SlZAR1 | Tomato ITAG release 2.40 (https://solgenomics.net/) | Solyc02g084890 | Materials and Methods: Bioinformatic and phylogenetic analyses |
|  |  | NCBI (https://www.ncbi.nlm.nih.gov/) | XP_004232433.1 |  |
| Gene (*Manihot esculenta*) | MeZAR1 | Manihot esculenta v7.1 (https://phytozome-next.jgi.doe.gov/) | Manes.02G097400 | Materials and Methods: Bioinformatic and phylogenetic analyses |
|  |  | NCBI (https://www.ncbi.nlm.nih.gov/) | P_021604862.1 |  |
| Gene (*Beta vulgaris*) | BvZAR1 | RefBeet-1.2 (http://bvseq.molgen.mpg.de/index.shtml) | Bv9_203570_geqf.t1 | Materials and Methods: Bioinformatic and phylogenetic analyses |
|  |  | NCBI (https://www.ncbi.nlm.nih.gov/) | XP_010688776.1 |  |
| Recombinant DNA reagent | ZAR1 plasmid collection | This paper | pZA1-52 | Materials and Methods: Plasmid constructions |

### Plant growth conditions

Wild type and mutant *N. benthamiana* were propagated in a glasshouse and, for most experiments, were grown in a controlled growth chamber with temperature 22-25°C, humidity 45-65% and 16/8-h light/dark cycle.

### Bioinformatic and phylogenetic analyses

We used NLR-parser (Steuernagel et al., 2015) to identify NLR sequences from the protein databases of tomato (Sol Genomics Network, Tomato ITAG release 2.40), *N. benthamiana* (Sol Genomics Network, *N. benthamiana* Genome v0.4.4), Arabidopsis (https://www.araport.org/, Araport11), cassava (https://phytozome-next.jgi.doe.gov/, Phytozome v13, Manihot esculenta v7.1) and sugar beet (http://bvseq.molgen.mpg.de/index.shtml, RefBeet-1.2). The obtained NLR sequences from NLR-parser, were aligned using MAFFT v. 7 (Katoh and Standley, 2013), and the protein sequences that lacked the p-loop motif were discarded to make the NLR dataset (Figure 1—source data 1). The gaps in the alignments were deleted manually in MEGA7 (Kumar et al., 2016) and the NB-ARC domains were used for generating phylogenetic trees (Figure 1—source data 2). The neighbour-joining tree was made using MEGA7 with JTT model and bootstrap values based on 100 iterations (Figure 1—source data 3). Amino acid sequences of ZAR1 orthologs and the percent identity are listed in Figure 1—figure supplement 1—source data 1 and —source data 2.

To calculate the phylogenetic (patristic) distance, we used Python script based on DendroPy (Sukumaran and Mark, 2010). We calculated patristic distances from each CC-NLR to the other CC-NLRs on the phylogenetic tree (Figure 1—source data 3) and extracted the distance between CC-NLRs of Arabidopsis or *N. bentha*miana to the closest NLR from the other plant species. The script used for the patristic distance calculation is available from GitHub (https://github.com/slt666666/Phylogenetic_distance_plot).

### Plasmid constructions

For Golden Gate cloning system, wild-type and MHD mutant clones of AtZAR1 and NbZAR1 were synthesized through GENEWIZ Standard Gene Synthesis with synonymous mutations to *Bpi*I restriction enzyme sites. To generate MADA motif mutants of AtZAR1 and NbZAR1, the leucine (L) 17 in the MADA motif was substituted to glutamic acid (E) by site-directed mutagenesis using Phusion High-Fidelity DNA Polymerase (Thermo Fisher). To make AtZAR1 and NbZAR1 clones for C-terminal tagging, the stop codons were removed by PCR using Phusion High-Fidelity DNA Polymerase (Thermo Fisher). Primers used for the site-directed mutagenesis and removing stop codon were listed in Supplementary file 1. All wild-type and mutant cording sequences were cloned into pCR8/GW/D-TOPO (Invitrogen) as level 0 modules (Weber et al., 2011), namely pZA1-16 (Table 1). The level 0 plasmids were then used for Golden Gate assembly with or without C-terminal tag modules, pICSL50009 (6xHA, Addgene no. 50309), pICSL50010 (4xMyc, Addgene no. 50310) or pICSL50034 (eGFP, TSL SynBio) into the binary vector pICH86988 (Engler et al., 2014), namely pZA17-52 (Table 1).

### Transient gene-expression and cell death assays

Transient expression of ZAR1 wild-type and mutants in *N. benthamiana* were performed by agroinfiltration according to methods described by Bos et al. (2006). Briefly, four-weeks old *N. benthamiana* plants were infiltrated with *A. tumefaciens* strains carrying the binary expression plasmids. *A. tumefaciens* suspensions were prepared in infiltration buffer (10 mM MES, 10 mM MgCl_2_, and 150 μM acetosyringone, pH5.6) and were adjusted to OD_600_ = 0.5. Macroscopic cell death phenotypes were scored according to the scale of Segretin et al. (2014) modified to range from 0 (no visible necrosis) to 7 (fully confluent necrosis).

### Protein immunoblotting

Protein samples were prepared from six discs (8 mm diameter) cut out of *N. benthamiana* leaves at 2 days after agroinfiltration and were homogenised in extraction buffer [10% glycerol, 25 mM Tris-HCl, pH 7.5, 1 mM EDTA, 150 mM NaCl, 2% (w/v) PVPP, 10 mM DTT, 1x protease inhibitor cocktail (SIGMA), 0.5% IGEPAL (SIGMA)]. The supernatant obtained after centrifugation at 12,000 x*g* for 10 min was used for SDS-PAGE. Immunoblotting was performed with HA-probe (F-7) HRP (Santa Cruz Biotech), c-Myc (9E10) HRP (Santa Cruz Biotech) or GFP (B-2) HRP (Santa Cruz Biotech) in a 1:5,000 dilution. Equal loading was checked by taking images of the stained PVDF membranes with Pierce Reversible Protein Stain Kit (#24585, Thermo Fisher).

### Plasmid distribution

All pZA plasmids will be available through Addgene.

## ACKNOWLEDGEMENTS

We are thankful to several colleagues for discussions and ideas. We thank Mark Youles of TSL SynBio for invaluable technical support. H.A. was funded by the Japan Society for the Promotion of Science (JSPS). This work was funded by the Gatsby Charitable Foundation, Biotechnology and Biological Sciences Research Council (BBSRC, UK), and European Research Council (ERC; NGRB and BLASTOFF projects).

## AUTHOR CONTRIBUTIONS

S.K. and H.A. designed the research, supervised the work and wrote the paper; A.H., T.S and H.A. performed research.

---

**Figure 1—figure supplement 1.**
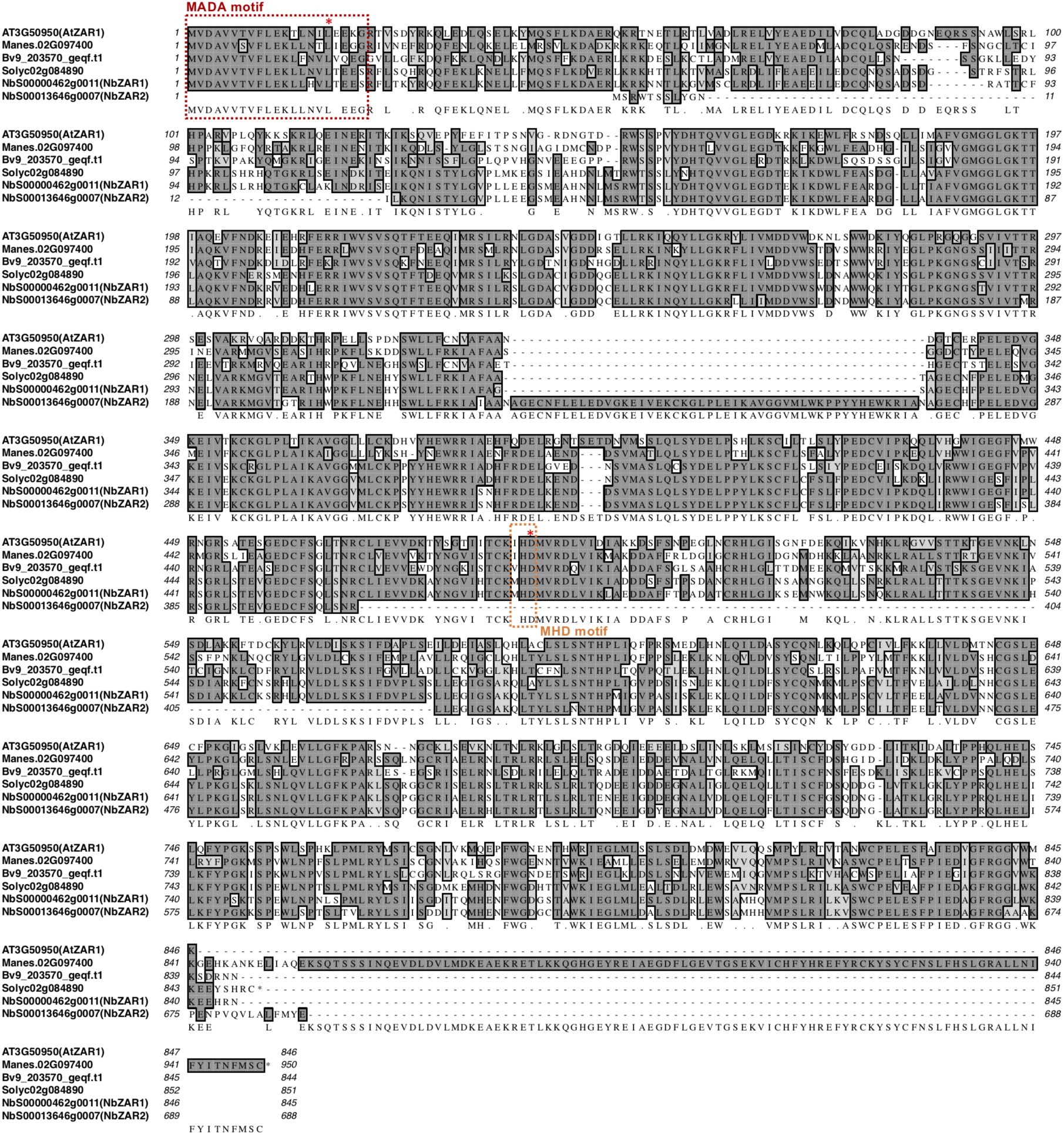
Sequence alignment of full-length ZAR1 ortholog proteins. Amino acid sequences of ZAR1 orthologs were aligned by ClustalW program. MADA and MHD motif sequences are marked with red and orange dot boxes, respectively. Red asterisks indicate substitution sites for MADA and MHD mutations.

**Figure 1—figure supplement 2.**
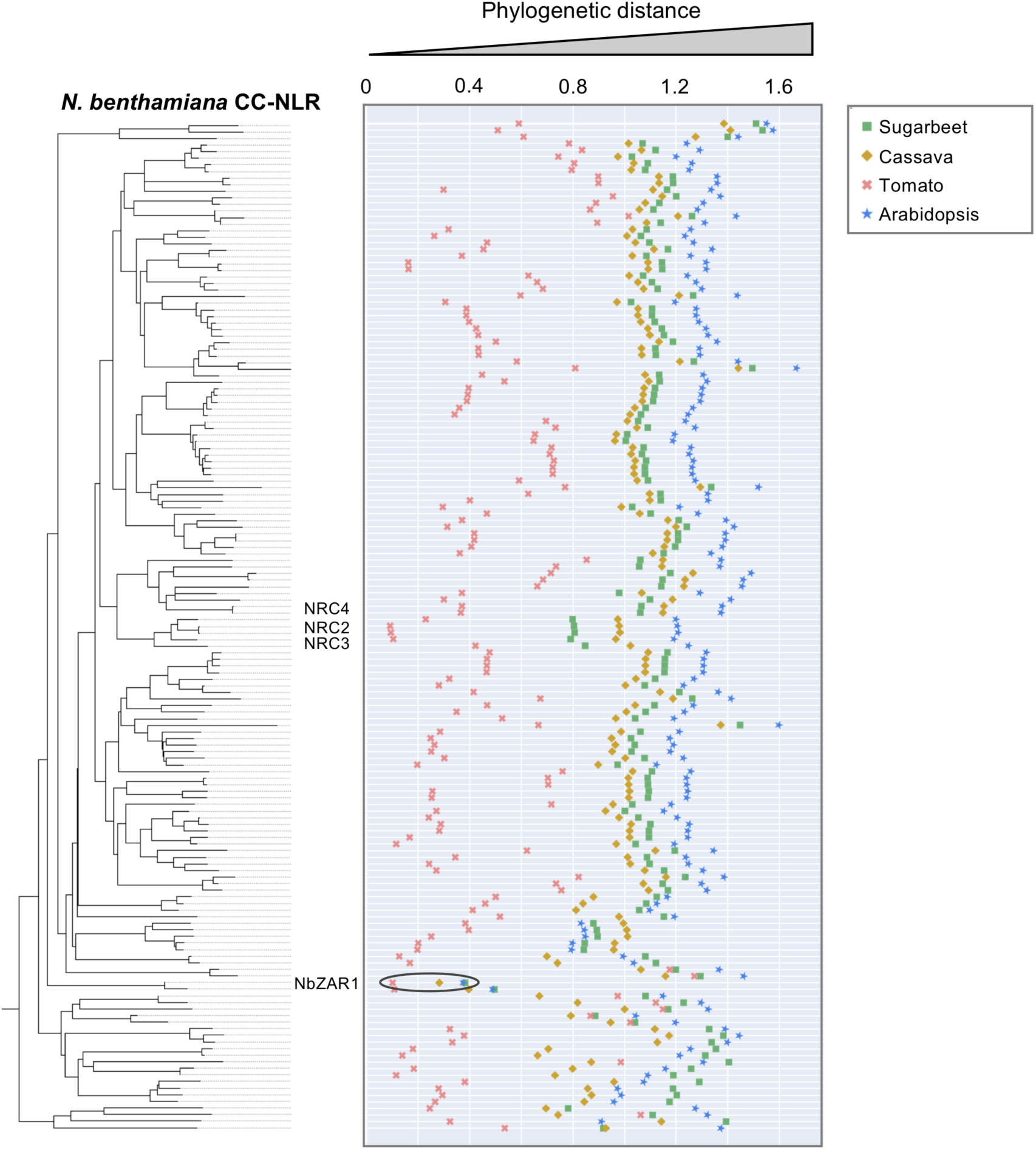
NbZAR1 is highly conserved across dicot species. The phylogenetic (patristic) distance of two CC-NLR nodes between *N. benthamiana* and the closest NLR from the other plant species were calculated from the NB-ARC phylogenetic tree in Figure 1A. The closest patristic distances are plotted with different colours based on plant species.

**Table 1—figure supplement 1.**
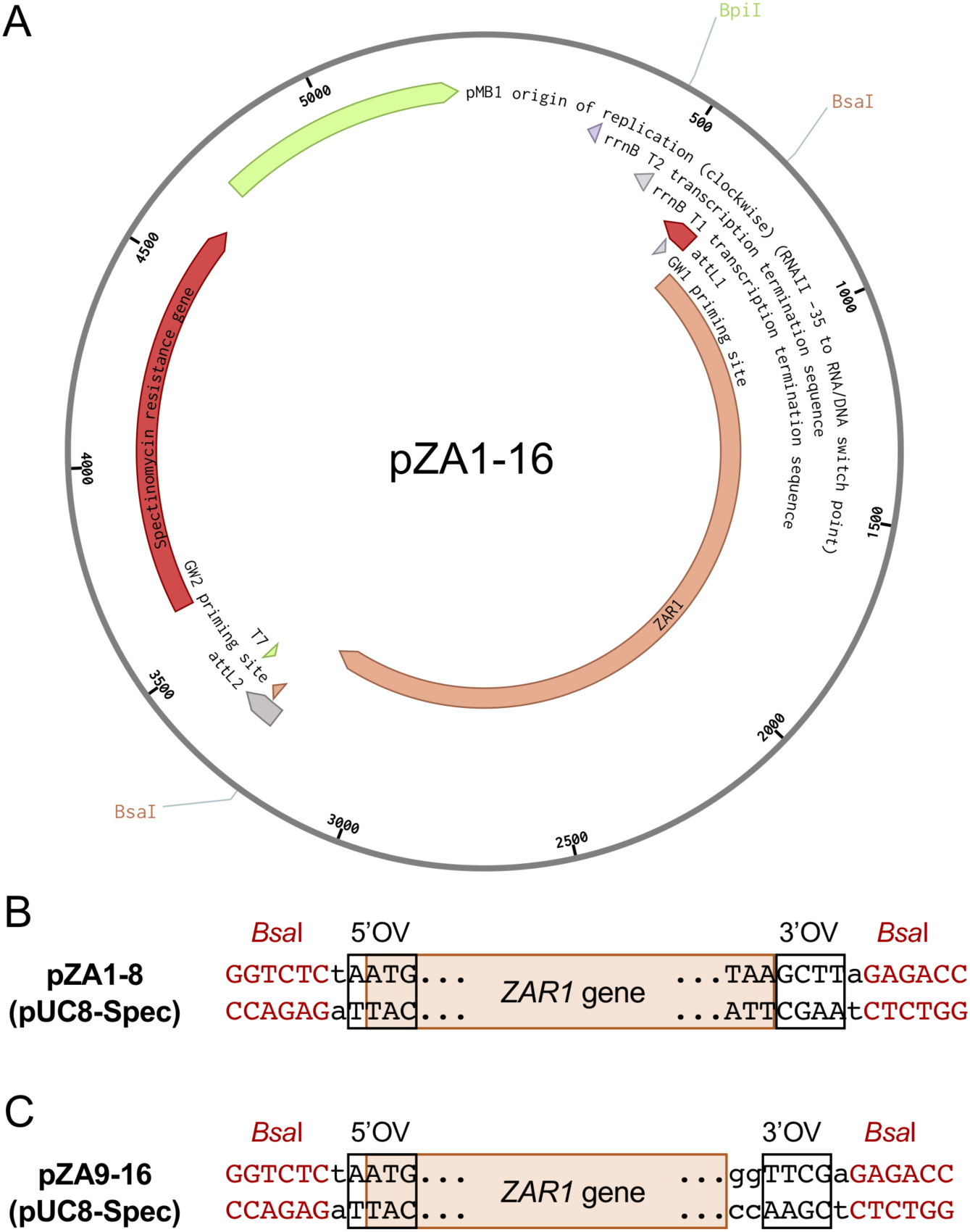
Golden Gate Level 0 modules of AtZAR1 and NbZAR1 genes. (**A**) AtZAR1 and NbZAR1 wild-type and the mutant genes were cloned in the pUC8-Spec vector (pZA1-16). The plasmid map was made with Benchling software. (**B**) ZAR1 genes were cloned with the stop codon in pZA1-8 plasmids. (**C**) *ZAR1* genes were cloned without the stop codon for C-terminal tag fusion (pZA9-16). *Bsa*I restriction enzyme sites for Golden Gate assembly are shown as red characters. 5’ and 3’ overhang sequences after *Bsa*I digestion are described as 5’OV and 3’OV, respectively.

## Supplementary files

**Supplementary file 1. Primers Used for generating ZAR1 Golden Gate Level 0 constructs**.

## Source data files

**Figure 1—source data 1. Amino acid sequences of full-length NLRs in the NLR dataset**. The 829 NLR sequences used in this study.

**Figure 1—source data 2. Amino acid sequences for NLR phylogenetic tree**. NB-ARC domain sequences used for phylogenetic analysis are shown with the IDs, *N. benthamiana* (NbS-), tomato (Solyc-), sugar beet (Bv-), Arabidopsis (AT-) and cassava (Manes-).

**Figure 1—source data 3. NLR phylogenetic tree file**. The phylogenetic tree was saved in newick file format.

**Figure 1**—**figure supplement 1—source data 1. Amino acid sequences of ZAR1 orthologs**. The N-terminal 28 amino acids (MAMRNQLYILESASQFPWAAFQKSEAEE) of SlZAR1 (Solyc02g084890) were removed due to potential mis-annotation.

**Figure 1**—**figure supplement 1—source data 2. Percent identity matrix of ZAR1 orthologs by Clustal2**.**1**.

