## Supplementary file 1 for "A vector system for fast-forward *in vivo* studies of the ZAR1 resistosome in the model plant *Nicotiana benthamiana*"

**Supplementary Table 1.** Primers Used for generating ZAR1 Golden Gate Level 0 constructs.

| Name | Sequence (5'>3') | Usage in this study |
|---|---|---|
| AtZAR1 BsaI Fw | AATGGTCTCTAATGGTGGACGCTGTTGTAAC | Level 0 construction |
| AtZAR1 STOP BsaI Rv | AATGGTCTCTAAGCTTAGGTTCTGTGCAATGG | Level 0 construction |
| AtZAR1 BsaI noSTOP Rv | AATGGTCTCTCGAACCGGTTCTGTGCAATGGTGTTTTC | Level 0 construction |
| AtZAR1 L17E BsaI Fw | AATGGTCTCTAATGGTGGACGCTGTTGTAACAGTGTTTTTA GAGAAAACCTTGAACATCgaaGAAGAAAAAGGCCGAACCG | MADA mutation |
| NbZAR1 BsaI Fw | AATGGTCTCTAATGGTGGATGCGGTGGTCAC | Level 0 construction |
| NbZAR1 STOP BsaI Rv | AATGGTCTCTAAGCTTAGTTCCTATGTTCTTCCTTC | Level 0 construction |
| NbZAR1 noSTOP BsaI Rv | AATGGTCTCTCGAACCGTTCCTATGTTCTTCCTTC | Level 0 construction |
| NbZAR1 L17E BsaI Fw | AATGGTCTCTAATGGTGGATGCGGTGGTCACTGTATTCTTG GAGAAACTTCTGCATGTTgaaACAGAGGAGAGTAGG | MADA mutation |
